# Enhancing the efficacy of siRNA Antibody Oligonucleotide Conjugates (AOCs) through chemical design

**DOI:** 10.64898/2026.09.23.753708

**Authors:** O. G. Hayes, J. Rädler, E. Filipiak, T. Czapik, S. Roudi, M. Ojansivu, H. Saranya Ilamathi, A. Marquant, Y. Huang, O. P.B. Wiklander, R. Zain, M. Honcharenko, S. EL Andaloussi

## Abstract

Antibody-oligonucleotide conjugates (AOCs) offer a promising solution to delivery challenges of therapeutic oligonucleotides. However, the relationship between their complex chemical architectures and biological activity remains poorly understood, limiting the development of important structure-function relationships. For siRNA-containing AOCs, increasing the siRNA-to-antibody ratio beyond one (drug-to-antibody ratio, DAR>1) reduces potency, attributed to altered pharmacokinetics arising from increased negative charge density. Here, we investigated whether simple chemical modifications could improve AOC efficacy and mitigate limitations associated with higher DAR. Introduction of a single C16 lipid modification to the siRNA significantly enhanced target gene silencing compared to the unmodified AOC. Extending this modification to a DAR2 architecture, in which two siRNAs are conjugated per antibody, restored the loss of activity associated with increasing DAR from 1 to 2. Notably, at an equivalent siRNA dose (1 mg/kg), the DAR2-C16 AOC requires half the amount of antibody while achieving knockdown comparable to the DAR1-C16 AOC. We also developed a charge-balancing ionizable linker (CBIL) designed to partially compensate for the negative charge of siRNA. Incorporation of the CBIL enhanced target gene silencing in heart and skeletal muscle without a corresponding increase in hepatic activity, resulting in a shift toward greater extrahepatic activity relative to liver. Together, these findings demonstrate that chemical modification of both the siRNA payload and antibody-siRNA linker can be used to tune AOC potency and tissue activity, while enabling higher payload loading without compromising efficacy. These results establish chemical design as an important strategy for expanding the architecture and therapeutic potential of AOCs.

**TOC Graphic:** 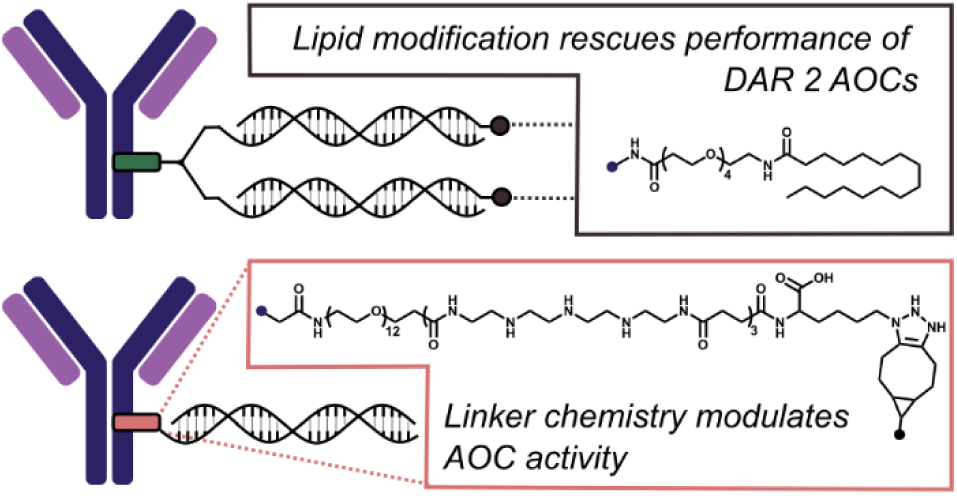

## Introduction

Therapeutic oligonucleotides treat disease by targeting the underlying genetic drivers of pathology, representing a highly versatile and powerful modality in modern drug development.^1,2^ For example, antisense oligonucleotides (ASOs) and small interfering RNAs (siRNAs) are short nucleic acid sequences typically 15-30 nucleotides in length that can selectively modulate gene expression through sequence-specific interactions with RNA transcripts. Despite this therapeutic promise, the clinical utility of such oligonucleotide therapeutics remains limited by inefficient delivery to specific tissue and cells, rendering much of the body “undruggable”.^3,4^

Oligonucleotides are inherently hydrophilic, negatively charged molecules that typically undergo rapid degradation and clearance *in vivo*. While chemical modifications of the phosphate backbone and 2’ position on the ribose ring have been developed to increase metabolic stability, targeted delivery of these molecules still presents a significant challenge.^5,6^ One of the most successful solutions to this problem is utilizing receptor-mediated delivery by conjugating oligonucleotides to molecules that drive recognition and delivery to cells in specific tissues.^7–10^ For example, conjugation of siRNA with multi-valent N-acetylgalactosamine (GalNAc) enables clathrin-dependent asialoglycoprotein (ASGP) receptor mediated endocytosis of siRNA for effective delivery to hepatocytes.^11^ Similarly, aptamer^12^ and peptide conjugates^13,14^ have been designed to target receptors of cells in specific organs and/or tumors for successful delivery of therapeutic oligonucleotides.

Antibodies are natural candidates for such receptor-targeted drug delivery mechanisms and the clinical success of antibody-drug conjugates (ADCs) in the delivery of chemotherapeutics to specific cells, while reducing toxic off-target effects, demonstrates their excellent suitability.^15,16^ Antibody oligonucleotide conjugates (AOCs) have emerged as a class of molecules that combine the sequence specificity of oligonucleotides and specific receptor affinity of antibodies, several of which are currently advancing through clinical development.^17–19^ Del-desiran, for example, is an AOC for the treatment of myotonic dystrophy type 1 (DM1) comprising an anti-transferrin receptor 1 antibody (αTfR1) and siRNA against DMPK mRNA. A recent phase I/II study reported effective delivery of del-desiran to muscle, with a mean ∼ 40% reduction in DMPK mRNA across all treated participants.^20^ Unfortunately, this substantial knockdown did not translate to a clinical benefit in Phase III, reflecting the challenges of treating complex diseases like DM1.^21^

While promising, AOCs are complex molecules and there is currently a limited understanding of how their vast chemical and architectural design space informs their therapeutic properties. Numerous interdependent design variables including antibody selection; conjugation strategy; linker chemistry; oligonucleotide chemistry; and drug-to-antibody ratio (DAR), can each influence pharmacokinetics, tissue distribution, intracellular trafficking, and ultimately efficacy. Consequently, establishing rational design principles for AOCs remains a significant challenge. One important example is DAR, where increasing the number of oligonucleotides per antibody beyond one has been reported to reduce the potency of both siRNA- and ASO-containing AOCs.^22,23^ This reduction has been attributed to increased overall negative charge, resulting in less favorable pharmacokinetics and enhanced systemic clearance. In contrast, antibody conjugates containing charge-neutral phosphorodiamidate morpholino oligonucleotides (PMOs) tolerate substantially higher DARs (up to 9.7) without compromising activity.^24^ Together, these observations suggest that overall molecular charge is an important determinant of AOC performance and highlight opportunities for improving therapeutic efficacy through rational chemical design. Developing strategies to effectively increase DAR without compromising performance has the potential to drastically reduce the quantity of antibody per therapeutic dose, increase potency by increasing number of siRNA molecules entering endosomal compartments and combine multiple, orthogonal, synergistic siRNA sequences into one architecture. To this end, recent work demonstrated that the incorporation of neutral backbone chemistries, such as phosphoryl guanidine, in siRNA led to improved *in vivo* performance of higher DAR AOCs.^25^ Although impactful, such chemistries require significant sequence optimization to avoid deleterious effects on intracellular activity.

Previous work from our group^26^ and others^27,28^ has demonstrated that lipid-conjugated oligonucleotides exhibit enhanced *in vivo* potency compared with their unmodified counterparts, primarily through favorable interactions with serum albumin that prolong circulation and increase tissue exposure.^29^ We therefore hypothesized that introducing a lipid modification (C16 palmitate) onto the siRNA component of an AOC could similarly enhance its therapeutic potency and potentially rescue the reduced efficacy observed at higher DARs. In parallel, we investigated whether reducing the overall negative charge of the conjugate through incorporation of a charge-balancing ionizable linker (CBIL) between the antibody and siRNA could further improve AOC performance. By evaluating these complementary chemical design strategies, this study seeks to establish simple, modular design principles for improving the potency and expanding the therapeutic potential of antibody oligonucleotide conjugates.

## Results and Discussion

### Selecting a model AOC system

To test the hypotheses outlined above, we focused our investigations on two readily modifiable features of the AOC architecture: the 3’ terminus of the sense strand and the linker connecting the siRNA and the antibody (Scheme 1a). To systematically evaluate the effects of these chemical design elements, we established a model AOC system in which all variables, other than the modifications under investigation, were held constant. Given the clinical validation of transferrin receptor 1 (TfR1/CD71) as a target for receptor-mediated oligonucleotide delivery,^30,31^ we selected an anti-mouse transferrin receptor 1 antibody (αmTfR1, 8D3) together with an siRNA that targets the murine superoxide dismutase (*Sod1*) gene. *Sod1* is broadly expressed across tissues, and this sequence has been validated in previous studies investigating siRNA conjugation chemistries.^26^ The full siRNA sequence as well as backbone and termini modifications are described in Scheme S1. Using this model system, we developed 6 AOC variants incorporating combinations of C16 lipid and a CBIL motif as well as structures with a defined DAR of 1 or 2 (Scheme 1b).

**Scheme 1.**
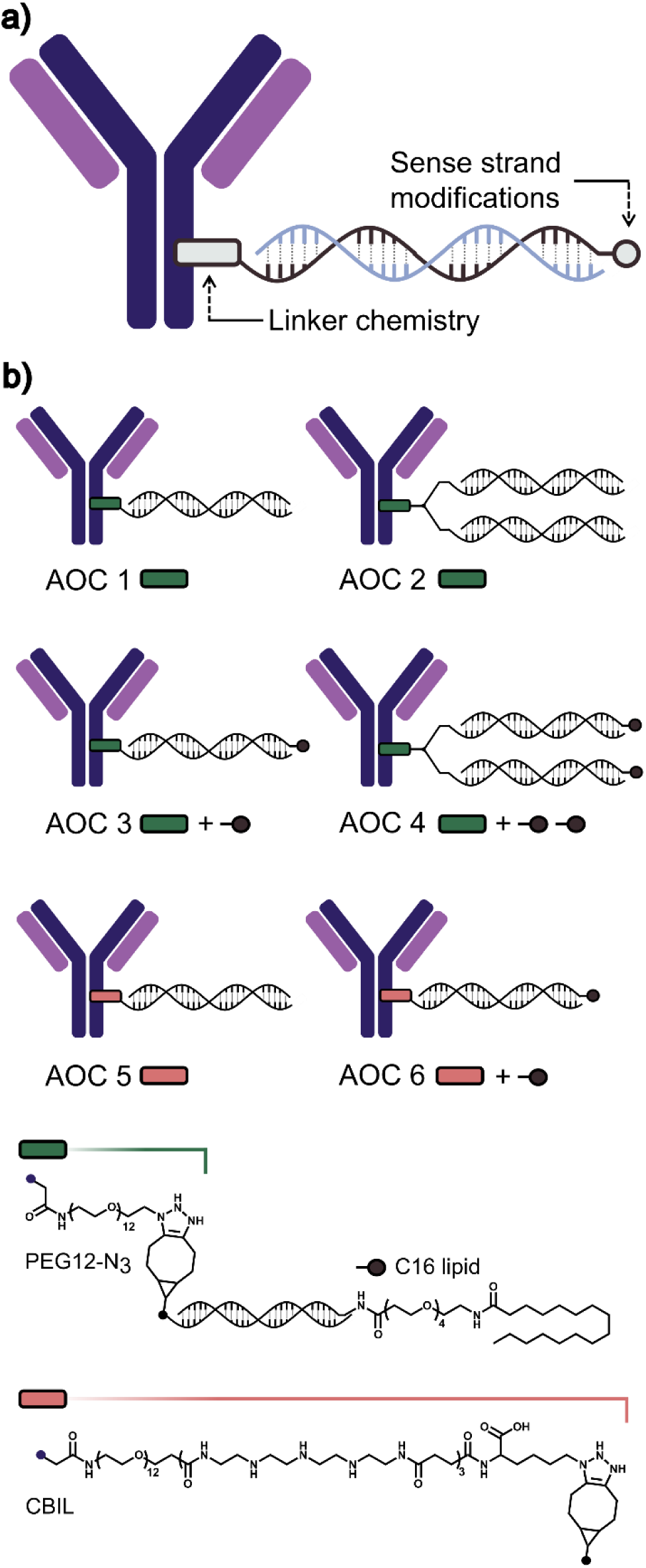
Overview of αmTfR1 AOC model system and the structural differences between AOCs. (a) Schematic representation of the AOC architecture and the sites of modification explored in this work. (b) Schemes of AOCs 1-6 where C16 lipid modification is represented by a black dot and the linkers, PEG12-N_3_ and CBIL, are represented by green and red boxes, respectively.

Numerous strategies have been reported for the preparation of AOCs,^32^ however, many rely on non-site-specific conjugation of siRNA to amino acid residues generating mixtures of positional isomers that can complicate interpretation of structure-function relationships.^33–35^ To mitigate the effects of siRNA conjugation location, we utilized a site-selective enzymatic strategy that enables installation of siRNA at an identical residue in the heavy chain (HC) across all constructs. This ensured homogenous AOC geometry and for differences in biological performance to be attributed primarily to the controlled variables rather than conjugation heterogeneity.

### Synthesis and characterization of AOCs

AOC variants were prepared using a universal two-step, site-selective conjugation workflow (Figure 1). First, the TfR1 antibody was enzymatically modified with a microbial transglutaminase (MTGase) to install a single azide functionality at a defined site (Q295) on each heavy chain. Conventional approaches typically perform deglycosylation with Peptide-N-Glyosidase F (PNGase F) to remove sterically hindering glycans from the HC followed by MTGase-mediated modification of Q295 in two separate steps. Here we developed a one-pot strategy in which deglycosylation and transglutamination were performed simultaneously (Figure 1a). Central to this method was the use of a previously reported MTGase variant with improved substrate scope and reactivity, enabling efficient modification of the HC with diverse linkers, such as the CBIL and PEG12-N_3_ linkers used in this work.^36^ Successful linker installation was confirmed by MS analysis (Figure 1b). Secondly, pre-annealed bicyclononyne (BCN) or dibenzocyclooctyne (DBCO) functionalized siRNA duplexes were conjugated to the antibody via strain-promoted azide–alkyne cycloaddition (SPAAC) chemistry, followed by removal of unconjugated siRNA by size-exclusion chromatography (SEC, Supporting Information 3.1.4). Importantly, pre annealed lipid-modified siRNAs and branched siRNAs (for DAR 2) could be readily incorporated without altering the overall synthetic workflow. This modular strategy enabled all AOC variants to be synthesized using an identical conjugation protocol, with structural diversity introduced solely through chemical modification of the siRNA and linker choice, therefore providing a straightforward and versatile platform for evaluating the effects of chemical design on AOC performance.

**Figure 1.**
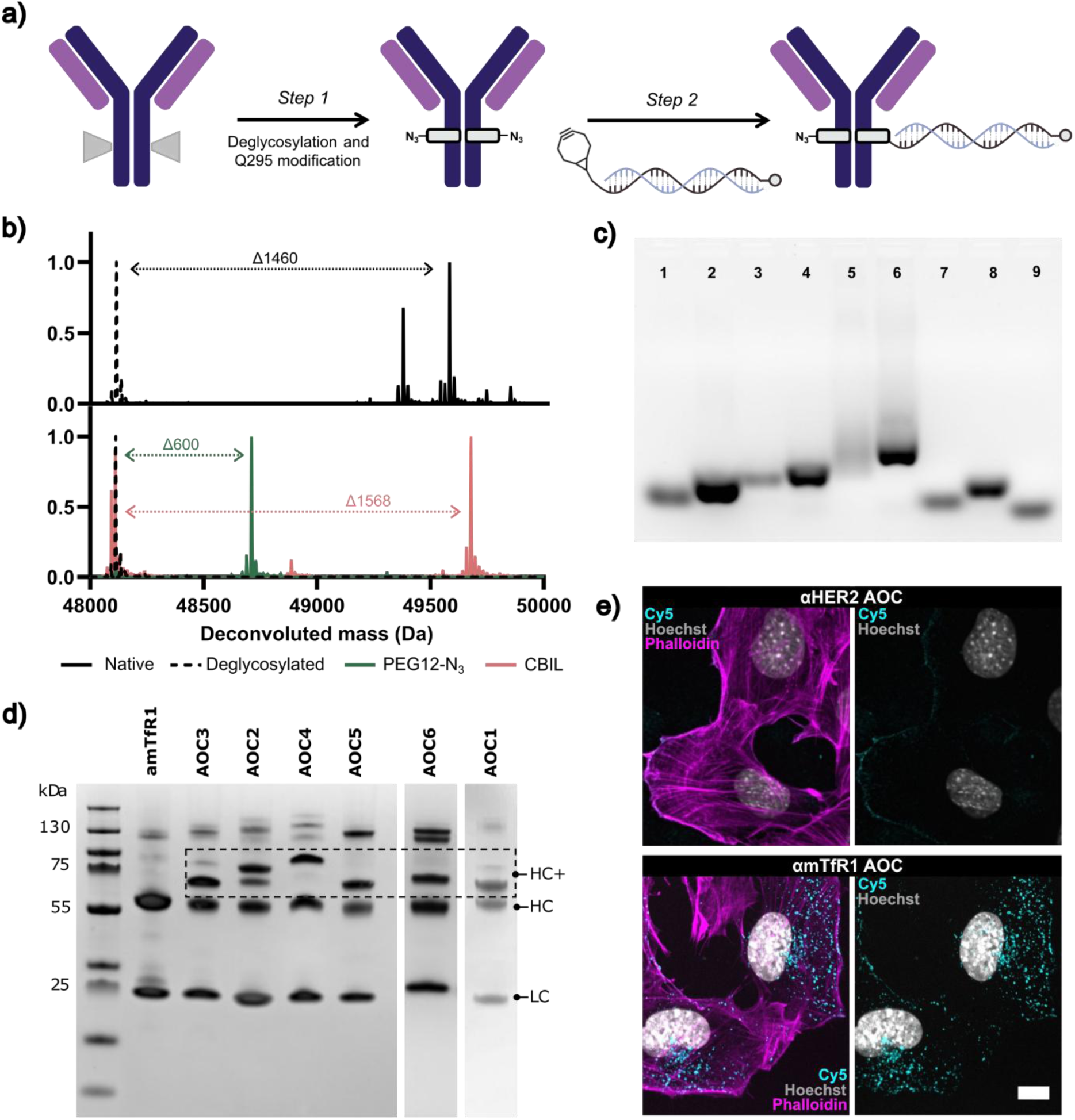
(a) Synthetic scheme showing the two-step synthesis protocol for preparation of AOCs. (b) Normalized, deconvoluted mass spec data revealing changes in mass of the heavy chain fragment after deglycosylation (top) and subsequent linker modification (bottom). (c) 2% native agarose gel of single stranded and duplexed oligonucleotides, visualized with GelRed. (1) S-C16, (2) S-C16 & AS, (3) branched-S, (4) branched-S & AS, (5) branched-S-C16, (6) branched-S-C16 & AS, (7) S, (8) S & AS, (9) AS, where S is sense strand and AS is antisense. (d) Denaturing SDS PAGE of AOCs, 4-12%, visualized using SimplyBlue stain. (e) C2C12 cells treated with αHER2 AOC control (top) or αmTfR1 AOC (bottom) were stained with phalloidin (magenta) to visualize F-actin and Hoechst (grey) to visualize nuclei. Left panels show merged F-actin, nuclear, and Cy5 fluorescence, while right panels show merged nuclear and Cy5 fluorescence, demonstrating cellular localization of the Cy5-labeled AOCs after 5 min incubation. Scale bar = 10 µm.

The C16 modified oligonucleotides incorporated into AOCs 3, 4 and 6, were synthesized according to a previously reported protocol (Supporting Information 2).^26^ In contrast, the branched Sod1 siRNAs used to generate DAR 2 architectures (AOCs 2 and 4) were developed specifically for this study. A detailed synthesis is provided in the Supporting Information; briefly, 5’ BCN bearing Sod1 sense strands (with or without 3’ C16 modification) were coupled to a tri-functional linker bearing two azide groups and one primary amine. After HPLC purification the remaining amine was reacted with an NHS-activated DBCO linker to generate the final click reactive branched siRNA. This design ensured that both siRNA duplexes were attached through a single branching point, providing a well-defined DAR 2 architecture while minimizing structural heterogeneity. Each synthetic intermediate was characterized by mass spectrometry (Supporting Information 2) and the final products were purified by HPLC, desalted and annealed with the corresponding Sod1 antisense strand prior to conjugation with antibodies. Native agarose gel electrophoresis analysis of all oligonucleotides confirmed efficient duplex formation and high purity across all constructs (Figure 1c).

Following conjugation, the resulting AOCs were characterized by SDS-PAGE to assess conjugation efficiency (Figure 1d). An unconjugated, deglycosylated αmTfR1 sample served as a reference for the migration of unmodified heavy and light chains. In all AOC samples, new bands with reduced electrophoretic mobility were observed above the heavy chain, consistent with successful siRNA conjugation. Across all constructs and reaction conditions (data not shown), the conjugation pattern consistently corresponded to approximately one successful click reaction per antibody. Therefore, although both heavy chains were expected to carry reactive azide groups following enzymatic modification (especially in the case of PEG12-N_3_ linker), only a single siRNA was typically conjugated to each antibody. We hypothesize that the first conjugation event proceeds rapidly, whereas attachment of a second negatively charged siRNA is kinetically disfavored owing to electrostatic repulsion, potentially compounded by steric effects. Densitometric analysis of SDS-PAGE profiles (Figure S10) further supports this showing comparable conjugation efficiencies across all six AOCs, ranging from 40.9 to 55.6% (overall mean of 48.7%). The similar conjugation efficiencies observed across the different constructs indicate that the chemical modifications to the siRNA and linker did not substantially affect antibody conjugation. The approximately 50% conjugation observed across constructs is consistent with predominantly mono-modified antibodies, suggesting that conjugation is largely restricted to a single heavy chain. Regardless of the underlying mechanism that dictates this behavior, it proves advantageous, as the overall DAR became defined by the siRNA architecture (single or branched) rather than the number of reactive sites on the antibody.

Furthermore, the substantial mass differences between the single and branched siRNA structures (∼8 kDa) produced the expected mobility shift, allowing AOCs containing branched siRNA (AOCs 2 and 4) to be readily distinguished from their single-siRNA counterparts. A lower molecular weight impurity was observed for AOC 2, corresponding to an antibody conjugated to a single siRNA. This impurity originated from incomplete purification of the branched siRNA precursor during HPLC and was determined unlikely to greatly influence the conclusions of the subsequent biological studies. Finally, no conjugation of the light chain was detected in any construct which is consistent with the site-selective modification strategy and the MS analysis of the linker modified antibodies (Figure S11).

Having established a robust synthetic workflow, we next sought to confirm that site-selective modification and siRNA conjugation did not compromise antibody-mediated cellular uptake. Murine C2C12 myoblasts were therefore incubated with two DAR 1 AOCs bearing a Cy5 fluorophore at the 3’ terminus of the siRNA sense strand: one prepared using αmTfR1 and a second prepared using a HER2 antibody, which served as a negative control since it binds human epidermal growth factor receptor 2 (HER2). Following a 5 min incubation, confocal microscopy revealed abundant intracellular Cy5-positive puncta in cells treated with the αmTfR1 AOC, whereas minimal fluorescence was observed for the trastuzumab control (Figure 1e). This punctate intracellular signal is consistent with rapid receptor-mediated uptake of the αmTfR1 AOC. Similar uptake profiles were observed following 30-minute incubations, with negligible uptake of the trastuzumab control throughout the experiment even after a 60-minute incubation (Figure S12). Together, these results demonstrate that the site-selective conjugation strategy preserves the receptor-targeting capability of the antibody, providing confidence that subsequent *in vitro* and *in vivo* studies reflect the influence of AOC chemical design rather than impaired antibody binding.

### Evaluating the impacts of chemical design of AOCs

Before evaluating the effects of chemical modifications on AOC performance, we first confirmed that the conjugated siRNA retained its RNAi activity and that antibody-mediated delivery resulted in productive intracellular gene silencing. Murine neuroblastoma (Neuro-2a, N2a) cells were treated with AOC 1 or unconjugated *Sod1* siRNA in the presence or absence of electroporation (Supporting Information 7). Inclusion of electroporated groups allowed assessment of the intrinsic activity of the siRNA independent of cellular delivery, while comparison of the non-electroporated groups evaluated the ability of the antibody to mediate receptor-dependent intracellular delivery.

As expected, electroporation of either unconjugated siRNA or AOC 1 resulted in robust *Sod1* mRNA knockdown (>95%), demonstrating that conjugation of the siRNA to the antibody resulted in retained silencing activity (Figure S13). In contrast, under non-electroporated conditions, unconjugated siRNA produced minimal knockdown (∼5%), whereas AOC 1 induced suppression of *Sod1* expression (∼26%) (Figure S14). These findings demonstrate that the AOC retains both the biological activity of the siRNA and the ability to mediate productive receptor-dependent intracellular delivery, providing a suitable platform for evaluating how subsequent chemical modifications influence AOC potency *in vivo*.

Guided by literature precedent, we performed a dose optimization trial to identify a dosage which would elicit sufficient knockdown of the target gene while also allowing differences to be observed when comparing AOCs. Female NMRI mice (n = 3 per group) were injected IV with AOC 1 at either 1 or 3 mg / kg, with respect to siRNA, and Sod1 mRNA levels were quantified in tissues following organ harvesting at day 7 post injection (Supporting Information 8.2). As transferrin receptor 1 is highly expressed in metabolically active tissues, we focused our analysis on heart and skeletal muscle, while including liver for comparison. A clear dose-dependent reduction in *Sod1* expression was observed across all three tissues (Figure S15). Based on these results, 1 mg/kg was selected for subsequence *in vivo* studies as it produced robust but incomplete knockdown. Importantly, both heart and skeletal muscle showed greater target knockdown than liver, consistent with preferential delivery mediated by TfR1 targeting.

We next investigated whether incorporation of a C16 lipid could enhance AOC efficacy and offset the reduced potency commonly associated with higher DAR constructs. To this end, lipid-modified and unmodified AOCs were prepared in both DAR 1 and DAR 2 formats (AOCs 1-4) and evaluated following IV administration at 1 mg / kg (with respect to siRNA, n = 5). Seven days after dosing, Sod1 mRNA levels were quantified in the liver, heart and skeletal muscle, and the activities of AOCs 1-4 were compared (Figure 2). As each treatment group received an identical siRNA dose, increasing the DAR from 1 to 2 was not expected to increase efficacy and based on previous reports, could potentially reduce potency. Consistent with this expectation, the non-lipidated DAR 2 construct (AOC 2) produced less *Sod1* knockdown than the corresponding DAR 1 construct (AOC 1), particularly in heart and skeletal muscle (Figure 2). Interestingly, this reduction in efficacy was accompanied by a marked change in the tissue profile of gene silencing. Whereas AOC 1 produced substantially greater knockdown in heart and skeletal muscle than in liver, AOC 2 generated comparatively similar levels of knockdown across all three tissues. Although mRNA knockdown alone cannot distinguish changes in tissue accumulation from differences in intracellular processing, these results indicate that increasing DAR substantially alters the *in vivo* performance of the AOC, consistent with previous reports that higher payload loading can negatively influence conjugate behavior.

**Figure 2.**
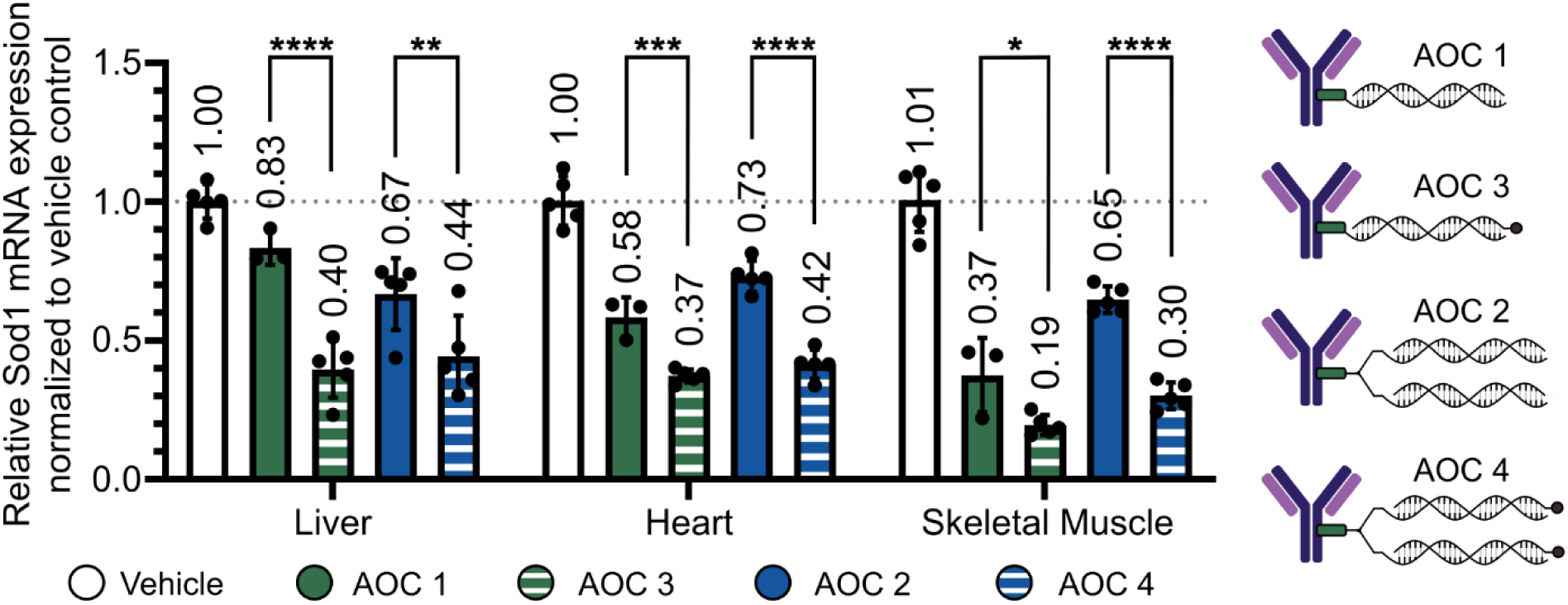
Investigating changes in target gene silencing activity for lipid modified AOCs. *Sod1* knockdown in liver, heart and skeletal muscle tissue normalized to vehicle control for AOCs 7 days post IV injection (1 mg/kg). Data are presented as mean ± SD of biological replicates (n = 3–5). Statistical significance was assessed separately for each organ using one-way ANOVA followed by Šídák’s multiple comparisons test. Comparisons shown represent the effect of C16 modification within each DAR group (AOC1 vs AOC3 and AOC2 vs AOC4). *P < 0.05, **P < 0.01, ***P < 0.001, ****P < 0.0001.

Comparison of the lipidated and non-lipidated AOCs revealed a pronounced improvement in efficacy following incorporation of the C16 modification (Figure 2). For both the DAR 1 and DAR 2 architectures, lipidated AOCs produced significantly greater *Sod1* knockdown than their corresponding non-lipidated counterparts across the tissues examined. Notably, the difference in activity between the lipidated DAR 1 (AOC 3) and lipidated DAR 2 (AOC 4) constructs was minimal, indicating that C16 modification largely mitigates the loss of potency associated with increasing DAR. Indeed, AOC 4 achieved target knockdown comparable to, and in some tissues greater than, the non-lipidated DAR 1 construct (AOC 1). Since all treatment groups received an equivalent siRNA dose, this finding suggests that similar therapeutic efficacy can be achieved using approximately half the quantity of antibody when employing a lipidated DAR 2 architecture. These results substantially expand the accessible design space of AOCs by demonstrating that higher DAR constructs can retain potent *in vivo* activity through appropriate chemical design. More broadly, increasing siRNA payload while maintaining efficacy may prove particularly advantageous for AOCs targeting receptors with limited expression or restricted rates of receptor-mediated internalization.

We next turned our attention to the development and evaluation of a charge-balancing ionizable linker (CBIL). Inspired by the widespread use of cationic polymers and ionizable lipids for the delivery of oligonucleotides in polyplexes and lipid nanoparticles (LNPs),^37,38^ respectively, we designed a polyamine-containing linker intended to partially compensate for the high negative charge associated with siRNA when incorporated into an AOC architecture. The CBIL was prepared by solid-phase peptide synthesis and comprised three units of Fmoc-TEPA(Boc3)-Suc, a C-terminal azidolysine residue, and an N-terminal PEG12-amine linker (Scheme 1, Supporting Information 3.1.2). Following deprotection, the resulting linker contains nine secondary aliphatic amines and therefore possesses a high density of ionizable sites. Structurally, the linker resembles a short, linear polyamine and, to some extent, mimics the charge characteristics of polyethylenimine (PEI).

Under physiological conditions, partial protonation of these amines is expected to generate a net positive charge, although the precise protonation state will depend on the local chemical environment and electrostatic interactions between neighbouring amines. If both heavy chains are modified, an antibody could therefore contain up to 18 ionizable amines, whereas modification of a single heavy chain would introduce 9. We hypothesized that protonation of these amines could partially screen the negative charge associated with the conjugated siRNA, thereby altering the overall physicochemical properties of the AOC and potentially mitigating the unfavorable effects associated with increasing anionic payload content. Notably, mass spectrometric analysis indicated a modestly lower extent of antibody modification with the CBIL compared with the PEG12-N3 linker (Figure 1b), which may reflect differences in the size, flexibility, or chemical properties of the two linker substrates. To further characterize the physicochemical consequences of CBIL incorporation, we analyzed the modified antibodies by ion-exchange chromatography (Figure S6). Antibodies modified with the CBIL exhibited increased retention compared with native and deglycosylated antibodies and those bearing the PEG12 linker, consistent with an increased positive surface charge arising from the polyamine-containing linker. Although ion-exchange chromatography does not directly recapitulate the physicochemical environment encountered *in vivo*, the altered retention provides evidence that incorporation of the CBIL substantially changes the physicochemical properties of the antibody.

We next evaluated whether the altered physicochemical properties imparted by the CBIL translated into changes in AOC activity *in vivo*. Comparison of *Sod1* mRNA knockdown following treatment with the CBIL-containing AOC 5 and the corresponding non-CBIL AOC 1 revealed enhanced activity in both heart and skeletal muscle, without a significant increase in hepatic knockdown (Figure 3a). The magnitude of the increase in extrahepatic knockdown was comparable to that observed following C16 modification in AOCs 3 and 4. However, unlike the lipidated AOCs, which exhibited increased knockdown in the heart, skeletal muscle and liver similarly, incorporation of the CBIL produced a more pronounced enhancement of extrahepatic activity relative to hepatic activity. These findings suggest that the two chemical modifications influence the tissue profile of AOC pharmacodynamic activity through distinct mechanisms and that charge-balancing may provide a means of preferentially enhancing extrahepatic activity. To further explore this phenomenon, we prepared an AOC that contained both the CBIL and a 3’ C16 lipid modification on the siRNA (AOC 6) and studied the knockdown of target mRNA *in vivo* (Figure 3b).

**Figure 3.**
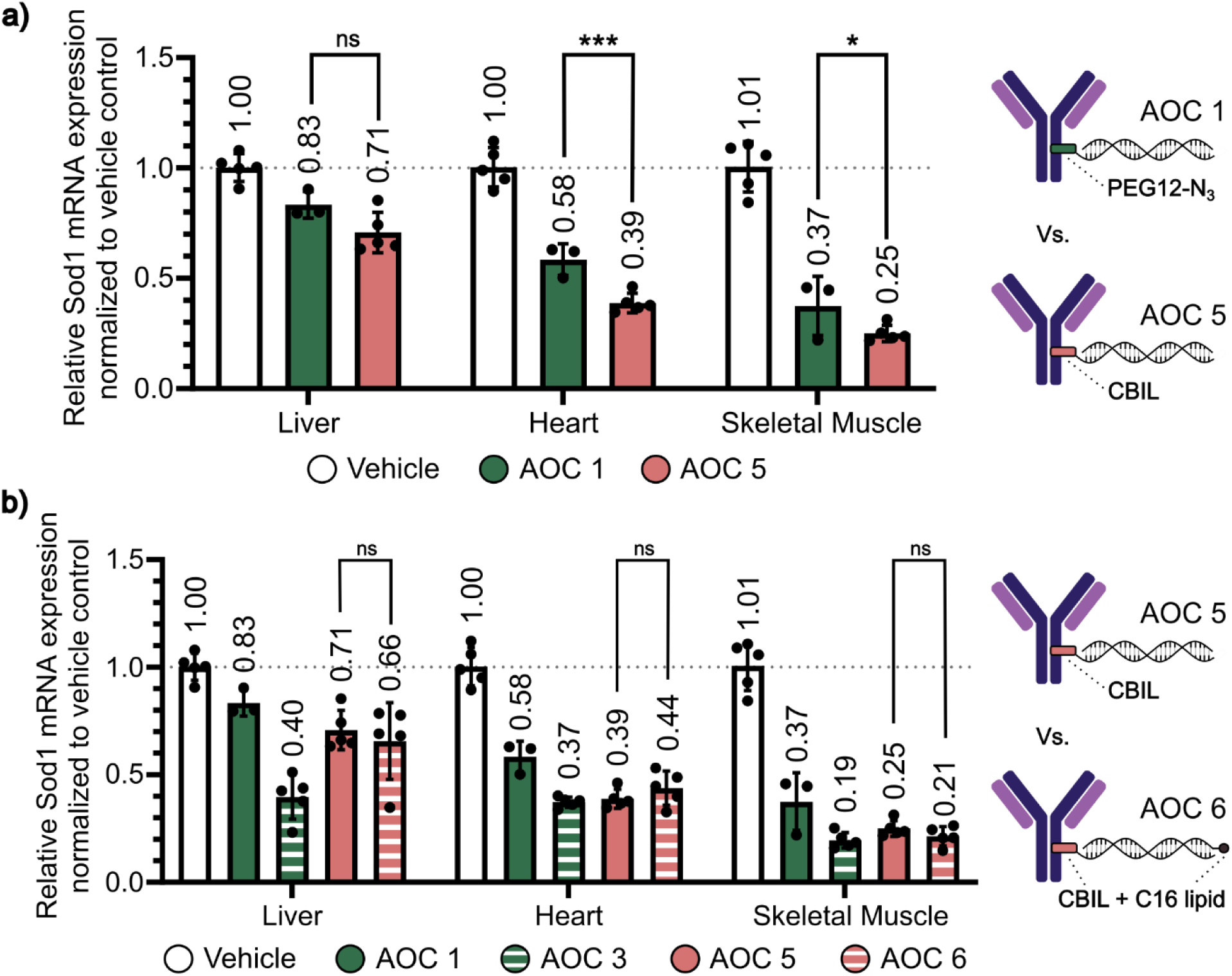
Investigating changes in target gene silencing activity for CBIL modified AOCs. (a) Comparison of target gene silencing between AOC 1 and 5. (b) Comparison of target gene silencing between AOC 5 and 6. *Sod1* knockdown in liver, heart and skeletal muscle tissue normalized to vehicle control for AOCs 7 days post IV injection (1 mg/kg). Data are presented as mean ± SD of biological replicates (n = 3–5). Statistical significance was assessed separately for each organ using one-way ANOVA followed by Šídák’s multiple comparisons test. *P < 0.05, **P < 0.01, ***P < 0.001, ****P < 0.0001.

Interestingly, the knockdown we observe for AOC 6 is not significantly different to that of AOC 5. Therefore, from this experiment, it is challenging to determine if any additional benefit is gained by having both the CBIL and C16 lipid in one structure. However, the absence of a significant difference does not establish that the effects of the two modifications are non-additive, as differences in pharmacodynamic kinetics or the magnitude and duration of tissue accumulation and silencing may not be captured at the single seven-day endpoint used in this study. A time-course analysis would therefore be required to determine whether the two architectures exhibit distinct temporal profiles of activity. Nonetheless, the persistence of reduced activity in the liver for AOC 6 in comparison to lipidated structures (AOC 3 & 4) suggests that incorporation of the CBIL attenuates the increase in hepatic activity while maintaining enhanced activity in extrahepatic tissues.

To further examine this tissue-selective activity profile, we calculated organ-to-liver knockdown ratios for each treatment group (Figure 4). AOCs containing C16 alone (AOCs 3 and 4) exhibited heart- or muscle-to-liver knockdown ratios of approximately 0.8-1.3, whereas CBIL-containing AOCs exhibited substantially higher ratios of approximately 1.6-2.6. Therefore, the CBIL-containing architectures showed a trend toward greater extrahepatic knockdown activity relative to hepatic activity. Although the mechanism underlying this difference remains unclear, these findings provide evidence that chemical modification of the AOC can influence not only overall potency but also the relative tissue profile of activity. Future studies incorporating direct pharmacokinetic and tissue biodistribution measurements, together with an expanded set of linker architectures, will be required to determine whether these effects arise from altered systemic exposure, tissue accumulation, intracellular trafficking, or combinations thereof.

**Figure 4.**
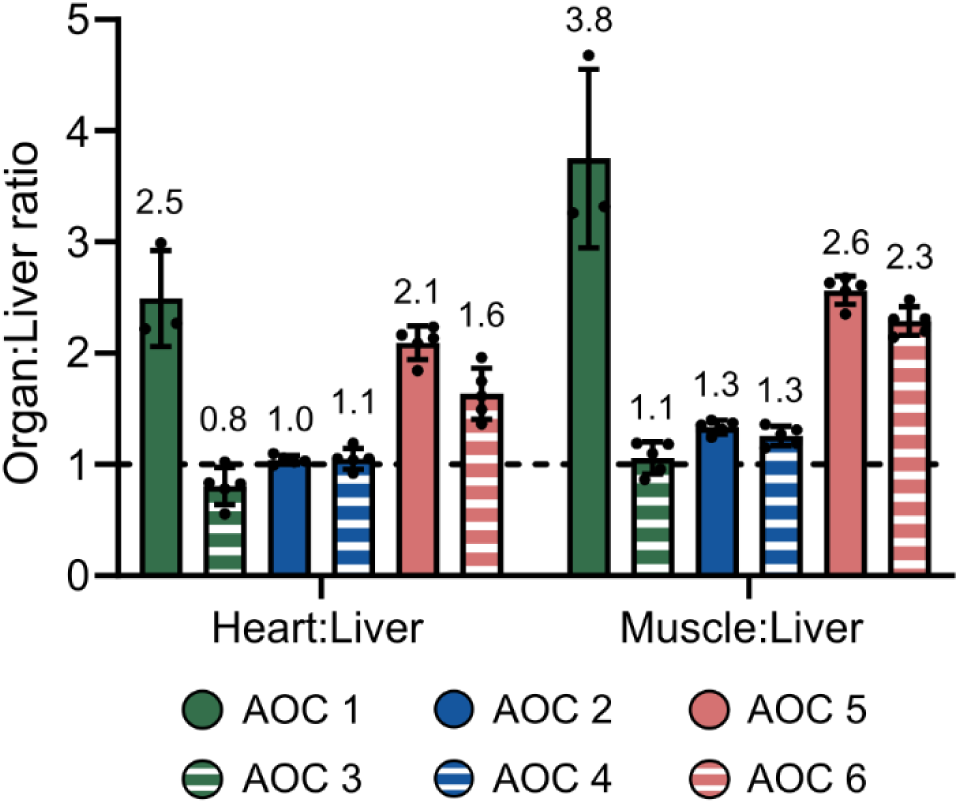
Organ-to-liver knockdown ratios for all AOCs. Each point represents an individual mouse, with the ratio calculated as the knockdown in the indicated organ divided by the mean liver knockdown of the corresponding treatment group. Bars indicate the mean ± S.

## Conclusions

This work explored how lipid modifications of siRNA and a novel antibody-siRNA linker can be used to increase the potency and expand the design space of AOC architectures. Specifically, we demonstrated that incorporation of a C16 palmitate modification, at the 3’ terminus of the sense strand, substantially enhanced *in vivo* AOC potency at both DAR 1 and DAR 2 in liver, skeletal muscle and heart tissues. Notably, the lipidated DAR 2 construct achieved target knockdown comparable to the DAR 1 AOC despite delivering the same siRNA dose with approximately half the number of antibody molecules. These findings demonstrate that appropriate chemical modification can mitigate the loss of potency associated with increased siRNA loading and thereby expand the accessible DAR space of AOCs. Such structures would be highly desirable when targeting receptors with low expression or when combining multiple, orthogonal siRNA sequences into a single AOC construct.

A charge-balancing ionizable linker containing multiple secondary amines was developed to investigate whether partial charge compensation could enhance AOC activity. We found that its incorporation altered the physicochemical properties of the antibody and enhanced *Sod1* knockdown in heart and skeletal muscle without a corresponding increase in hepatic activity. In contrast to C16 modification, which broadly enhanced activity across tissues, the CBIL produced a more pronounced shift toward extrahepatic pharmacodynamic activity relative to liver. Combining the CBIL and C16 modifications retained this enhanced extrahepatic activity, although no additional increase in overall potency was observed under the conditions examined.

Together, these results demonstrate that simple chemical modifications, including siRNA lipidation and linker design, can be used not only to increase AOC potency but also to modulate their tissue profile of pharmacodynamic activity. In particular, lipid modification provides a strategy for maintaining potency at higher DAR, while charge-balancing chemistry offers a complementary approach for enhancing extrahepatic activity relative to the liver. These findings highlight the potential of chemical design to expand the AOC design space beyond the choice of antibody and oligonucleotide cargo. The modular nature of AOCs provides numerous opportunities to independently tune properties such as payload loading, potency, and tissue activity through rational chemical design. Further studies combining pharmacokinetic, tissue distribution, and intracellular trafficking measurements will be important for establishing the mechanisms underlying these effects and developing generalizable design principles for next-generation AOCs.

## Supporting Information

Oligonucleotide structures, sequences and molecular weights; Synthesis and characterization methods of AOCs; Mass spectrometry data for all modified oligonucleotides; Ion exchange chromatography analysis; Comprehensive confocal data; *In vitro* evaluation of AOCs; Dose-optimization study *in vivo*; Further details regarding animal experiments.

## Authors

Julia Rädlar – Division of Biomolecular and Cellular Medicine, Department of Laboratory Medicine, Karolinska Institutet, 14152 Stockholm, Sweden; Department of Cellular Therapy and Allogeneic Stem Cell Transplantation (CAST), Karolinska University Hospital, 14186 Stockholm, Sweden.

Eliza Filipiak – Division of Biomolecular and Cellular Medicine, Department of Laboratory Medicine, Karolinska Institutet, 14152 Stockholm, Sweden; Department of Cellular Therapy and Allogeneic Stem Cell Transplantation (CAST), Karolinska University Hospital, 14186 Stockholm, Sweden.

Tomasz Czapik – Division of Biomolecular and Cellular Medicine, Department of Laboratory Medicine, Karolinska Institutet, 14152 Stockholm, Sweden; Department of Cellular Therapy and Allogeneic Stem Cell Transplantation (CAST), Karolinska University Hospital, 14186 Stockholm, Sweden.

Samantha Roudi – Division of Biomolecular and Cellular Medicine, Department of Laboratory Medicine, Karolinska Institutet, 14152 Stockholm, Sweden; Department of Cellular Therapy and Allogeneic Stem Cell Transplantation (CAST), Karolinska University Hospital, 14186 Stockholm, Sweden; Karolinska ATMP (Advanced Therapy Medicinal Products) Center, Karolinska Institutet, 14152 Stockholm, Sweden.

Miina Ojansivu – Division of Biomolecular and Cellular Medicine, Department of Laboratory Medicine, Karolinska Institutet, 14152 Stockholm, Sweden; Department of Cellular Therapy and

Allogeneic Stem Cell Transplantation (CAST), Karolinska University Hospital, 14186 Stockholm, Sweden. Karolinska ATMP (Advanced Therapy Medicinal Products) Center, Karolinska Institutet, 14152 Stockholm, Sweden.

Hema Saranya Ilamathi – Division of Biomolecular and Cellular Medicine, Department of Laboratory Medicine, Karolinska Institutet, 14152 Stockholm, Sweden; Breast Center, Karolinska Comprehensive Cancer Center, Karolinska University Hospital, 14186, Stockholm, Sweden Antonin Marquant – Division of Biomolecular and Cellular Medicine, Department of Laboratory Medicine, Karolinska Institutet, 14152 Stockholm, Sweden; Department of Cellular Therapy and Allogeneic Stem Cell Transplantation (CAST), Karolinska University Hospital, 14186 Stockholm, Sweden; Karolinska ATMP (Advanced Therapy Medicinal Products) Center, Karolinska Institutet, 14152 Stockholm, Sweden

Yanjie Huang – Division of Biomolecular and Cellular Medicine, Department of Laboratory Medicine, Karolinska Institutet, 14152 Stockholm, Sweden; Department of Cellular Therapy and Allogeneic Stem Cell Transplantation (CAST), Karolinska University Hospital, 14186 Stockholm, Sweden.

Oscar Wiklander – Division of Biomolecular and Cellular Medicine, Department of Laboratory Medicine, Karolinska Institutet, 14152 Stockholm, Sweden; Breast Center, Karolinska Comprehensive Cancer Center, Karolinska University Hospital, 14186, Stockholm, Sweden

Rula Zain – Division of Biomolecular and Cellular Medicine, Department of Laboratory Medicine, Karolinska Institutet, 14152 Stockholm, Sweden; Karolinska ATMP (Advanced Therapy Medicinal Products) Center, Karolinska Institutet, 14152 Stockholm, Sweden; Center for Rare Diseases, Clinical Genetics and Genomics, Karolinska University Hospital, SE-17176 Stockholm, Sweden

## Acknowledgements

Part of this work was facilitated by the Protein Science Facility at Karolinska Institutet, Stockholm, (and we would like to thank Dr Emilia Strandback, Dr Tom Reichenbach, Dr Henrik Spåhr and Dr Tomas Nyman for assistance). Also, imaging for this study was performed at the Luminous Cell Imaging core facility/Nikon Center of Excellence, at Karolinska Institutet, Sweden, supported by the KI infrastructure board (1–48/2024) and the Olle Engvist Foundation (235-0496).

