## Supporting Information for "Enhancing the efficacy of siRNA Antibody Oligonucleotide Conjugates (AOCs) through chemical design"

### Supporting Information for
### Enhancing the efficacy of siRNA Antibody Oligonucleotide Conjugates (AOCs) through chemical design

*O. G. Hayes†‡*, J. Rädler†‡, E. Filipiak†‡, T. Czapik†‡, S. Roudi†‡§, M. Ojansivu†‡§, H. Saranya Ilamathi†#, A. Marquant†‡§, Y. Huang†‡, O. P.B. Wiklander†#, R. Zain†‡§⊥, M. Honcharenko†‡§* and S. EL Andaloussi‡†§**

**Author Addresses**

†Division of Biomolecular and Cellular Medicine, Department of Laboratory Medicine, Karolinska Institutet, Huddinge, 14152, Stockholm, Sweden.

‡Department of Cellular Therapy and Allogeneic Stem Cell Transplantation (CAST), Karolinska University Hospital, 14186 Stockholm, Sweden.

§Karolinska ATMP (Advanced Therapy Medicinal Products) Center, Karolinska Institutet, 14152 Stockholm, Sweden

#Breast Center, Karolinska Comprehensive Cancer Center, Karolinska University Hospital, 14186, Stockholm, Sweden

⊥Center for Rare Diseases, Clinical Genetics and Genomics, Karolinska University Hospital, SE-17176, Stockholm, Sweden

*Corresponding authors

**1. List of abbreviations**

ACN - Acetonitrile

AS – Antisense strand

BCN – Bicyclononyne

DCBO – Dibenzocylooctyne

DIC - Diisopropylcarbodiimide

DMF - Dimethylformamide

DMSO – Dimethyl sulfoxide

FPLC – Fast protein liquid chromatography

HPLC – High performance liquid chromatography

LC-MS – Liquid chromatography mass spectrometry

NHS - N-Hydroxysuccinimide

NMP – N-Methyl-2-pyrrolidone

OD – Optical density

PBS – Phosphate buffered saline

PEG – Polyethylene glycol

SS – Sense strand

TEAA – Triethyl ammonium acetate

**2. Structure and synthesis of oligonucleotides**

2.1 General structure of chemically modified Sod1 siRNA

Oligonucleotides used in this study were acquired from Axolabs GmbH and then modified accordingly. The sequences and backbone chemical modifications are described in scheme S1.

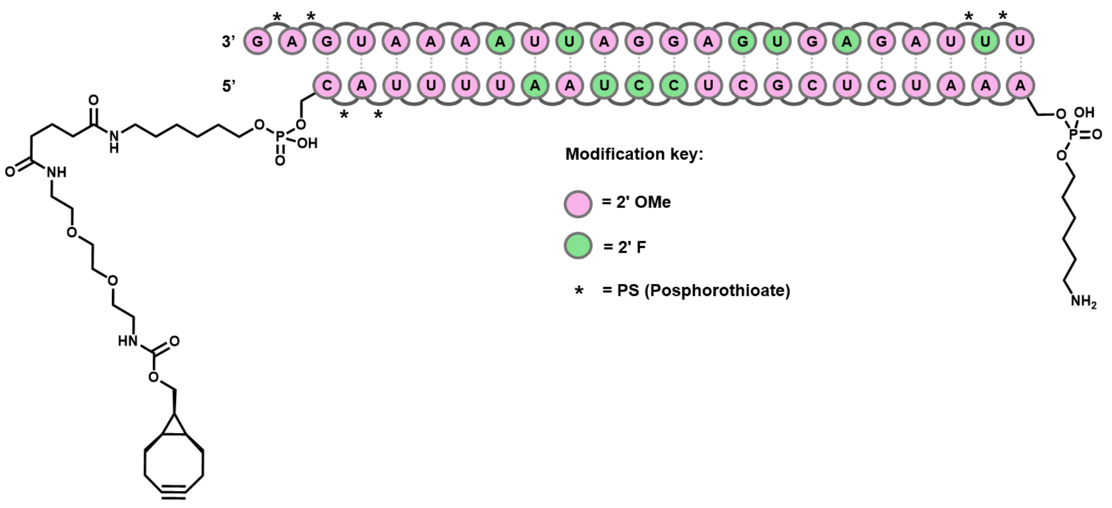

**Scheme S1.** Schematic representation of *Sod1* siRNA structure using comprising antisense (AS) and sense strand B (SSB).

#### 2.2 C16-lipid modification

In a 2 mL eppendorf, 2 equiv. of palmitic acid-PEG4-NHS ester (BroadPharm, DMSO) was added to 1 equiv. of SSB dissolved in pH 8.8 carbonate buffer. The reaction was incubated at 37 degrees with agitation on a table top shaker for 1 hour. Excess palmitic acid was removed by extraction with 3x washes of ethylacetate. The reaction was then purified using RP-HPLC (A= 10% ACN in 50 mM TEAA buffer, B= 100% ACN). Fractions containing SSB-peg4-C16 were collected and freeze-dried. LC-MS analysis of SSB-peg4-C16 confirmed the mass.

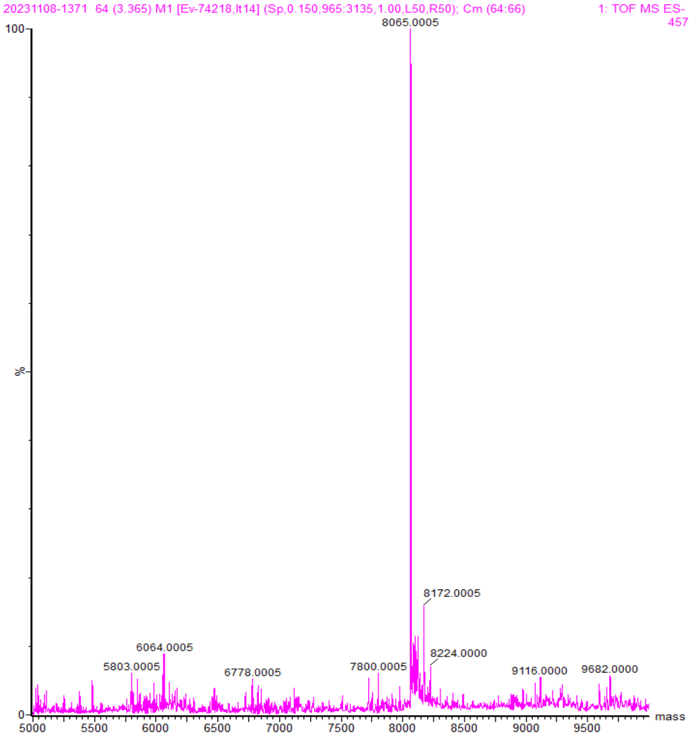

**Figure S1.** Deconvoluted mass spectrum of SSB-C16

2.3 Synthesis of branched siRNA

In a 2mL eppendorf, 2.5 equiv. of SSA or SSB-peg4-C16 (90% $H_2O$, 10% ACN solution) was added to 1 equiv. of N-(Amino-PEG10)-N-bis(PEG10-azide) (BroadPharm BP-25640, DMF). The reaction was incubated at ambient temperature with agitation on a tabletop shaker overnight. The reaction was then purified using RP-HPLC (A= 10% ACN in 50 mM TEAA buffer, B= 100% ACN), and fractions containing product were collected and freeze dried. LC-MS analysis confirmed formation of branched oligos.

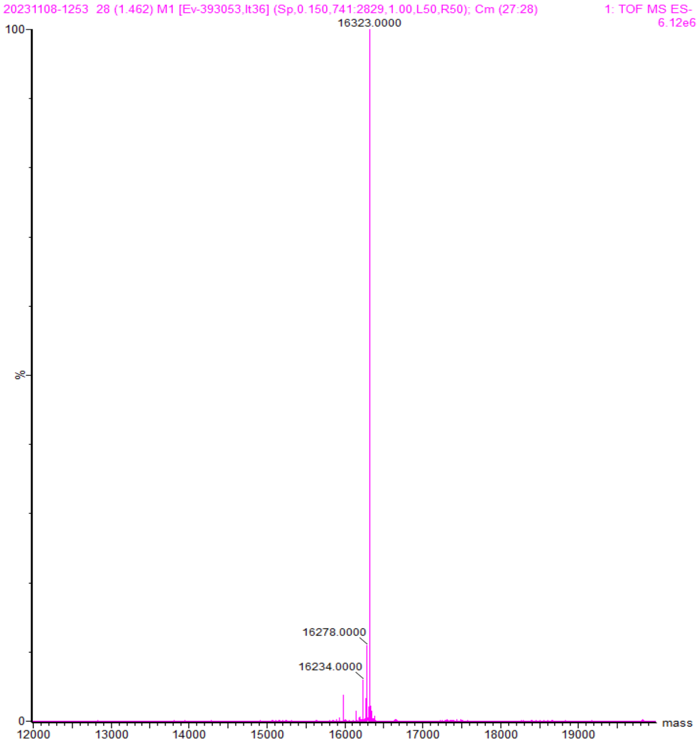

**Figure S2.** Deconvoluted mass spectrum of branched-SSA

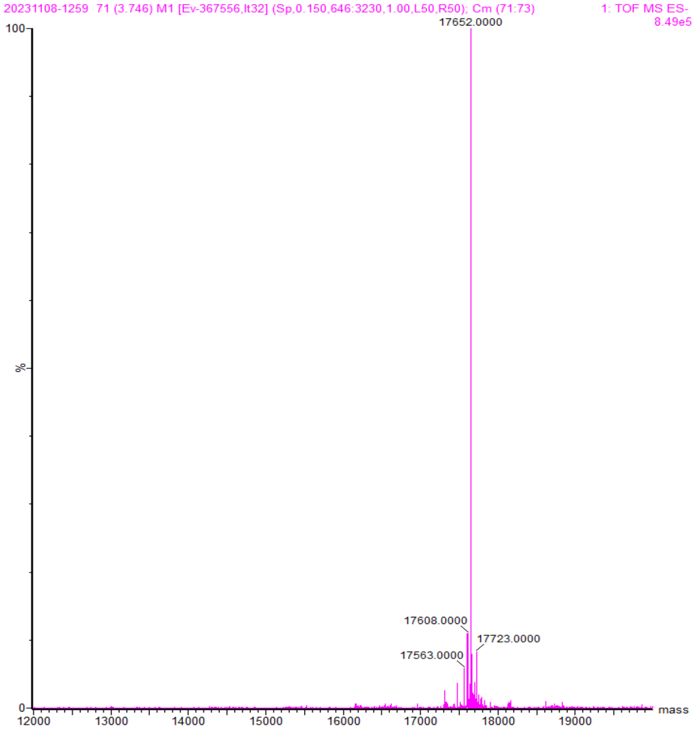

**Figure S3.** Deconvoluted mass spectrum of branched-SSB-C16

Next, 1 equiv. of branched oligo (dissolved in pH 8.8 carbonate buffer) was modified with 1.2 equiv. DBCO-NHS ester (Lumiprobe). The reaction proceeded for 1 hr at ambient temperature, with agitation on the tabletop shaker. The reaction was then quenched with 1M Tris buffer and unreacted small molecules were removed using NAP10 Sephadex columns (Cytiva). Final products were analyzed by LC-MS and freeze-dried before use.

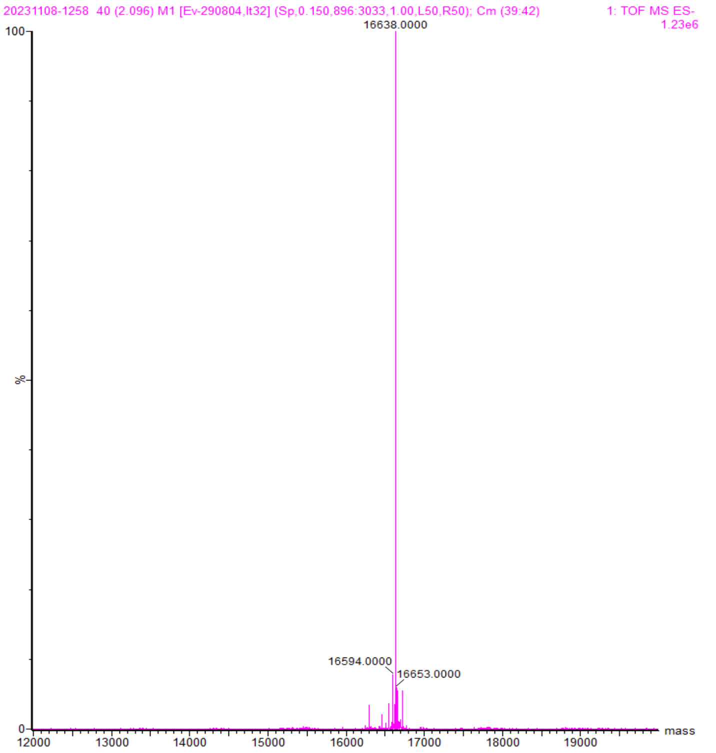

**Figure S4.** Deconvoluted mass spectrum of DBCO-branched-SSA

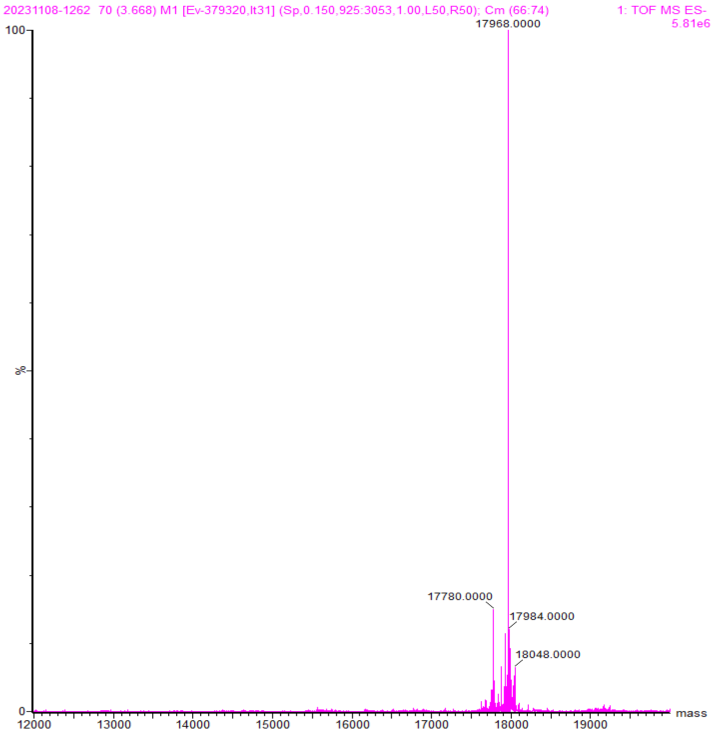

**Figure S5.** Deconvoluted mass spectrum of DBCO-branched-SSB-C16

2.4 Synthesis of Cy5-labelled siRNA

In a 2 mL eppendorf, 2 equiv. of Sulfo-Cy5-NHS ester (Lumiprobe, DMSO) was added to 1 equiv. of SSB dissolved in pH 8.8 carbonate buffer. The reaction was incubated at 25 degrees with agitation on a tabletop shaker for 1 hour. The reaction was then purified using RP-HPLC (A= 10% ACN in 50 mM TEAA buffer, B= 100% ACN). Fractions containing product were collected and freeze-dried. LC-MS analysis of SSB-Cy5 confirmed the mass.

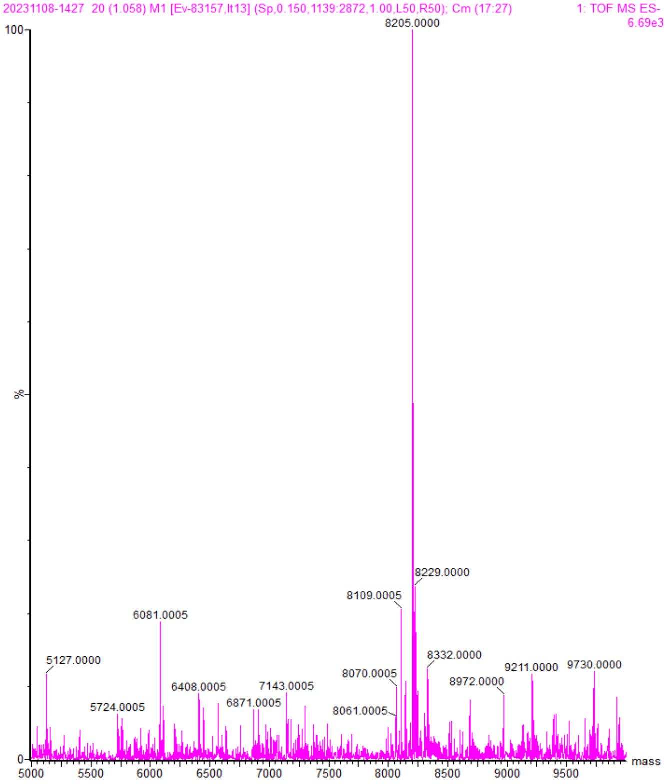

**Figure S6.** Deconvoluted mass spectrum of SSB-Cy5

2.4 Table of oligonucleotides and molecular weights

| Name | Sequence (5′ to 3′) | Calculated Mw (Da) | Found Mw (Da) |
|---|---|---|---|
| SSA (Sense Strand A) | BCN-CAUUUUAAUCCUCACUCUAAA | 7401.1 | 7399.9 |
| SSB (Sense Strand B) | BCN-CAUUUUAAUCCUCACUCUAAA-NH2 | 7580.3 | 7579.4 |
| AS (Antisense) | UUUAGAGUGAGGAUUAAAAUGAG | 7775.4 | 7774.4 |
| SSB-C16 | BCN-CAUUUUAAUCCUCACUCUAAA-peg4-C16 | 8066.1 | 8065.0 |
| DBCO-branched-SSA | DBCO-(CAUUUUAAUCCUCACUCUAAA)$_2$ | 16637.1 | 16638.0 |
| DBCO-branched-SSB-C16 | DBCO-(CAUUUUAAUCCUCACUCUAAA-peg4-C16)$_2$ | 17967.1 | 17968.0 |

| | | | |
|---|---|---|---|
| SSB-Cy5 | BCN-CAUUUUAAUCCUCACUCUAAA-Cy5 | 8204.2 | 8205.0 |

**3. Synthesis and characterization of AOCs**

3.1 MTGase-mediated modification of Ab with azide-linker

In a 1.5 mL eppendorf tube, 250 µl of anti-mouse CD71 (TfR1, bioXcell BE0329, 4.8 mg/mL), 2µl of PNGase F (500 units/ µl, New England Biolabs), 50 ul of mMTGase (1x PBS, 5mg/mL) and 1 ul of azido-peg12-amine (200mg/mL, DMSO, BroadPharm BP-29725) were mixed in a total volume of 400 µl (made up with 1xPBS) and incubated at 37 °C overnight. Subsequently, purification of the modified antibody was performed by size exclusion chromatography, using an Äkta Pure instrument (Cytiva) equipped with a Superdex 200 increase 10/300 GL size exclusion column (Cytiva). Mobile phase: 1x PBS and peak detection from absorbance at 280 nm. Fractions were collected and concentrated in an Amicon spin filter (100kda MWCO). For modification of the Ab with CBIL, identical conditions were used but 3 ul of a 200 mg/mL linker solution (DMF) was added.

3.1.1 Mutant MTGase (mMTGase)

mTGase sequence (containing N-terminal pro-peptide)

MDNGAGEETKSYAETYRLTADDVANINALNESAPAASSAGPSFRAPDSDDRVTPPAEPLDRMPDP
YRPSYGRAETVVNNYIRKWQQVYSHRDGRKQQMTEEQREWLSYGCVGVTWVNSGQYPTNRLA
FASFDEDRFKNELKNGRPRSGETRAEFEGRVAKESFDEEKGFQRAREVASVMNRALENAHDESAY
LDNLKKELANGNDALRNEDARSPFYSALRNTPSFKERNGGNHDPSRMKAVIYSKHFWSGQDRSS
SADKRKYGDPDAFRPAPGTGLVDMSRDRNIPRSPTSPGESFVNFDYGWFGAQTEADADKTVWTH
GNHYHAPNGSLGAMHVYESKFRNWSDGYSDFDRGAYVITFIPKSWNTAPDKVKQGWPLEAHHH
HHH*

The construct was transformed into E. coli BL21 (DE3) STAR pRARE2. The cells were cultivated in Terrific Broth (TB) medium and incubated at 37°C with shaking (175 rpm). At different times, the OD was measured for the culture and the temperature was set to 18°C at OD 2. The protein expression was induced at approximately OD 3 (IPTG, final concentration 0.1 mM). Protein expression continued overnight before the cells were harvested by centrifugation (10 min at 4500 × g). The protein was then purified from cell lysate by immobilized metal ion chromatography (IMAC), followed by size exclusion chromatography (SEC).

mMTGase was activated by proteolytic cleavage with 0.5 U/mL dispase I (Sigma Aldrich) for 30 min at 37 °C. Cleaved mMTGase was purified by an additional IMAC purification step and then buffer-exchanged to MTG buffer (25 mM Tris HCl, 150 mM NaCl, pH 8.0) for direct use or storage at -80 °C.

3.1.2 Synthetis of charge-balancing-ionizable-linker (CBIL)

Peptide-based linker was carried out on Biotage Initiator microwave peptide synthesizer using Fmoc chemistry and DIC\Oxyma as a coupling agents under the nitrogen gas. Rink Amide ChemMatrix resin (213 mg, 100 µmol) was placed in 10 mL reactor vial and swollen in NMP for 20 min in 70°C. Deprotection of Fmoc groups was performed by using 20% piperidine in NMP for 13 min at room temperature. Peptide couplings were conducted by using 5 eq of amino acid monomers (Fmoc-TEPA(Boc3)-Suc, Amino-PEG12-acid), 5 eq of DIC and 5 eq of Oxyma in NMP for 6 min in 75°C. Capping step was performed by using NMP-lutidine-acetic anhydrate (89:6:5 v\v\v) for 2 min at room temperature. After completion of the synthesis, the resin was washed with NMP and DCM. Peptide was deprotected and cleaved from solid support by mixing the resin with 10 mL of a deprotection mixture of TFA-TIS-$H_2O$ (95:2.5:2.5 v\v\v) for 2 h at room temperature. The resin was filtrated, and the filtrate was directly drained into 30 mL of cold methyl tert-butyl ether. The crude precipitate was separated by centrifugation, then re-dissolved in 10 mL of $H_2O$-acetonitrile (50:50 v\v) and freeze-dried. Crude peptide was purified by RP-HPLC on a Waters XBridge Prep C18 5 µm OBD 19x100mm column with 10 mL\min flow rate using linear gradient elution 10-95 % of Acetonitrile with 0.2% TFA in $H_2O$ with 0.2% TFA in 16 min using detection at 265 nm and 280 nm. LC-MS confirmed analysis of CBIL confirmed the mass.

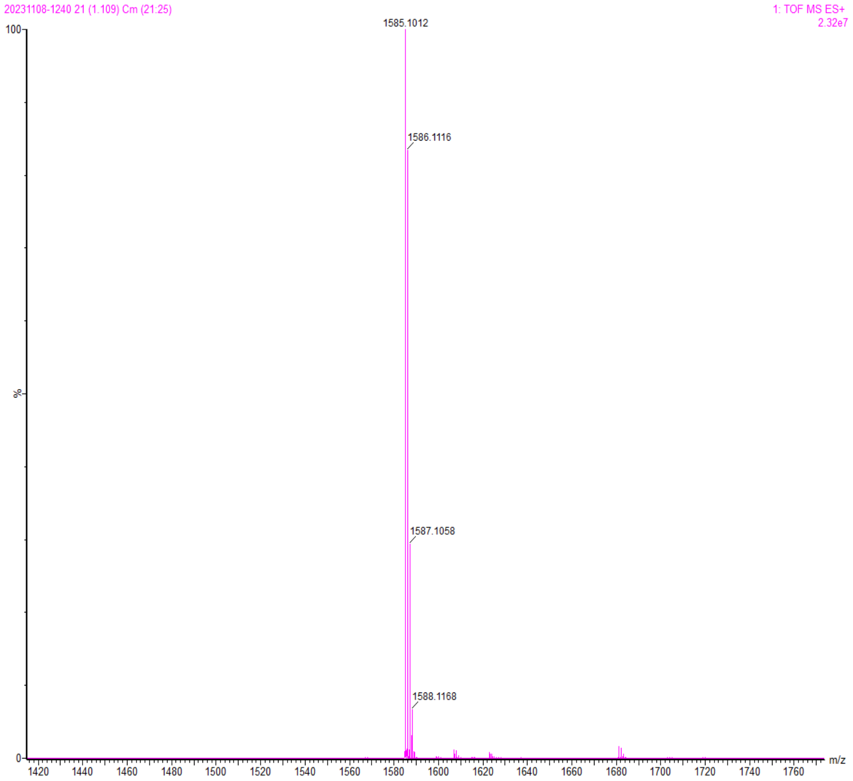

**Figure S7.** Mass spectrum of CBIL.

3.1.3 Ion-exchange chromatography analysis of linker modified Abs

Antibodies were analyzed by cation-exchange chromatography using a 1 mL HiTrap Capto S ImpAct column (Cytiva) connected to an Äkta FPLC system. Buffer A consisted of 50 mM sodium acetate, pH 5.5, and Buffer B consisted of 50 mM sodium acetate, pH 5.5, containing 1 M NaCl. The column was equilibrated with Buffer A prior to sample injection. Following sample loading, the column was washed with Buffer A and bound antibody species were eluted using a linear gradient of Buffer B. Protein elution was monitored by UV absorbance at 280 nm, and chromatograms were analyzed based on retention volume and peak profile.

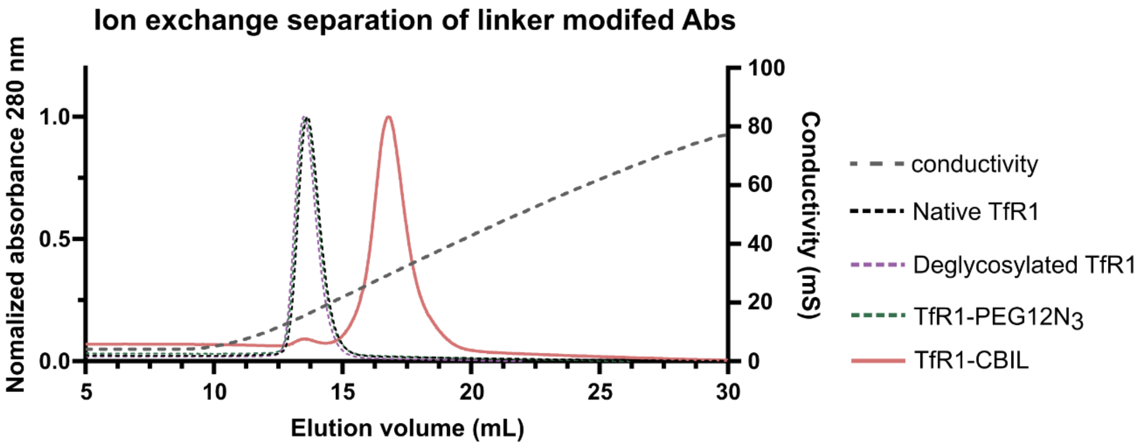

**Figure S8.** Ion exchange chromatograms of amTfR1 and its derivatives.

3.1.4 Size Exclusion Chromatography (SEC)

Antibodies and antibody-oligonucleotide conjugates (AOCs) were purified by size-exclusion chromatography (SEC) using a Superdex™ 200 Increase 10/300 GL column (Cytiva) connected to an ÄKTA™ go chromatography system. The column was equilibrated with at least 2 column volumes (CV) of 1× PBS prior to sample application. Samples were injected at a volume of ≤500 µL. The sample was applied to the column and separated at a flow rate of 0.75 mL/min at room temperature, with absorbance monitored at 280 nm. Fractions corresponding to the antibody/AOC peak were collected and analyzed by SDS-PAGE. Fractions containing the desired product were pooled and concentrated using centrifugal filters with a 100 kDa molecular weight cutoff. The column was equilibrated with 1× PBS between consecutive runs.

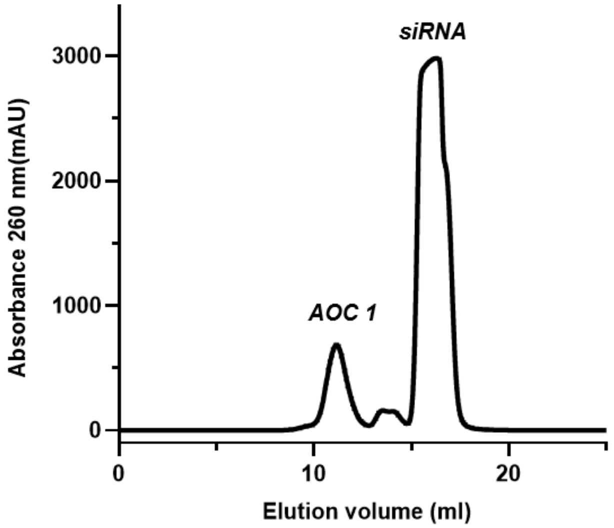

**Figure S9.** Representative SEC chromatogram from purification of AOC 1 from excess siRNA.

3.2 siRNA conjugation and characterization

Prior to conjugation, BCN-modified sense and anti-sense strands were annealed at 300 µM by heating to 95 °C for 10 min, followed by incubation at 37 °C for 60 min. 25 equiv. of duplex siRNA was then added to azide-modified CD71 and incubated at room temperature overnight with gentle agitation. Subsequently, purification of the antibody-oligonucleotide conjugate was performed by size-exclusion chromatography to remove unconjugated siRNA. Fractions collected and concentrated in amicon spin filker (100kda MWCO). siRNA may be modified with lipids (C16) or branched linkers prior to conjugation reaction.

##### 3.2.1 Densiometry analysis

SDS-PAGE gels were analyzed by densitometry using ImageJ. A region of interest was drawn around the heavy chain bands in each antibody lane (Figure SXa), and the intensity profile was generated using the Plot Profile function (Figure SXb). The unconjugated and conjugated antibody bands were analysed by integration of corresponding peaks. Conjugation efficiency was calculated as the integrated intensity of the conjugated species divided by the total integrated intensity of conjugated and unconjugated species, expressed as a percentage (Figure SXc).

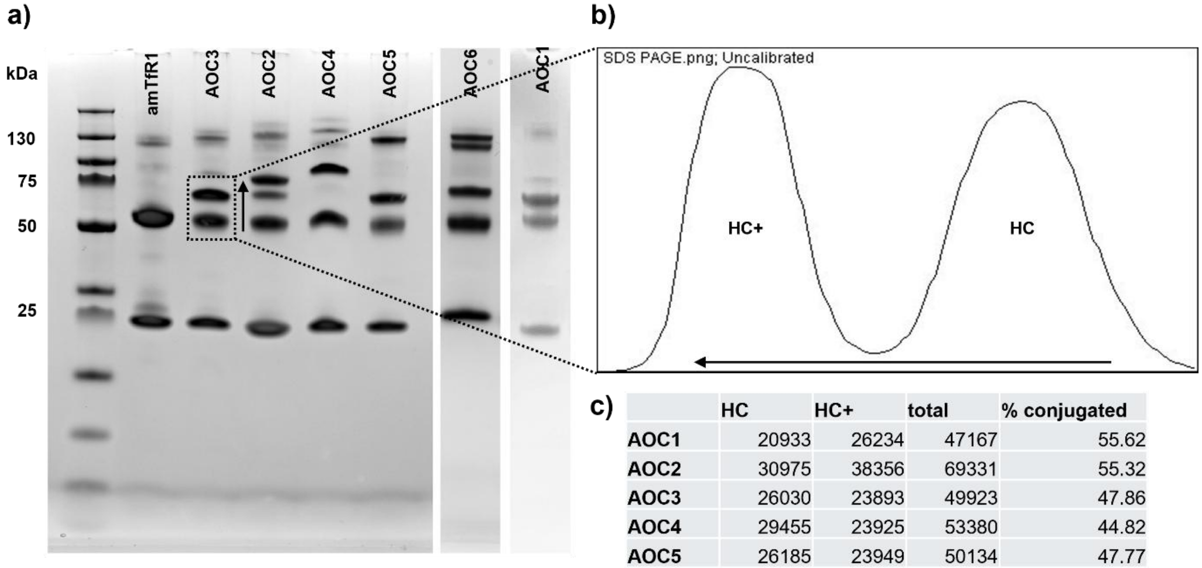

c)

| | HC | HC+ | total | % conjugated |
|---|---|---|---|---|
| **AOC1** | 20933 | 26234 | 47167 | 55.62 |
| **AOC2** | 30975 | 38356 | 69331 | 55.32 |
| **AOC3** | 26030 | 23893 | 49923 | 47.86 |
| **AOC4** | 29455 | 23925 | 53380 | 44.82 |
| **AOC5** | 26185 | 23949 | 50134 | 47.77 |
| **AOC6** | 35217 | 24359 | 59576 | 40.89 |

**Figure S10.** Densitometric analysis of antibody-siRNA conjugation. (a) SDS-PAGE gel used to assess antibody–siRNA conjugation. (b) Representative intensity profile generated from the indicated lane using the Plot Profile function in ImageJ, showing the peaks corresponding to unconjugated and conjugated antibody species. (c) Integrated intensities of the corresponding bands and calculated conjugation efficiencies for the six AOCs. Conjugation efficiency was calculated as the integrated intensity of the conjugated species relative to the total integrated intensity of conjugated and unconjugated species.

##### 3.2.3 AOC concentration determination

The concentration of siRNA-containing samples was determined using the QuantiFluor® RNA System (Promega) and Quantus™ Fluorometer according to the manufacturer's instructions. Briefly, QuantiFluor RNA Dye was diluted 1:400 in 1× TE buffer to prepare the working solution for high-concentration samples. For each measurement, 200 µL of dye working solution was added to a 0.5 mL PCR tube, followed by 1–20 µL of sample. Samples were mixed thoroughly and incubated for 5 min at room temperature, protected from light. The Quantus Fluorometer was calibrated using the provided RNA standard (500 ng) and a blank containing dye working solution prior to measurement. Sample concentrations were determined using the RNA measurement mode of the instrument, accounting for the volume of sample added. Samples were diluted in nuclease-free water or 1× TE buffer as required to fall within the assay's quantification range.

#### 4. LC-MS methods

4.1 LC-MS analysis of oligonucleotides

An ACQUITY UPLC BEH C18 (1.7 µm) 2.1 x 50 mm column (p/n 186002350) was employed to analyze conjugated oligonucleotides using mobile phase of water (A) and acetonitrile (B), both containing 5 mM ammonium acetate. Samples were injected (10 µl) onto the column (heated to 60 °C) and separation was achieved using a flowrate of 0.8 mL/min and a gradient of 10 to 90% B over 8 minutes. Mass detection was performed using the Xevo™ G2XS QToF mass spectrometer in negative ionization mode.

4.2 LC-MS analysis of antibodies

A BioResolve RP mAb Polyphenyl column (450 Å, 2.7 µm, 2.1 x 150 mm) was employed to analyze antibodies using a mobile phase of (A) water and (B) acetonitrile, both containing 0.1% difluoroacetic acid. Samples were injected (10 µL) onto the column (heated to 80 °C) and separation was achieved using a flowrate of 0.2 mL/min and a gradient of 10 to 50% B over 6 minutes. Mass detection was performed using the Xevo G2XS QToF mass spectrometer in positive ionization mode.

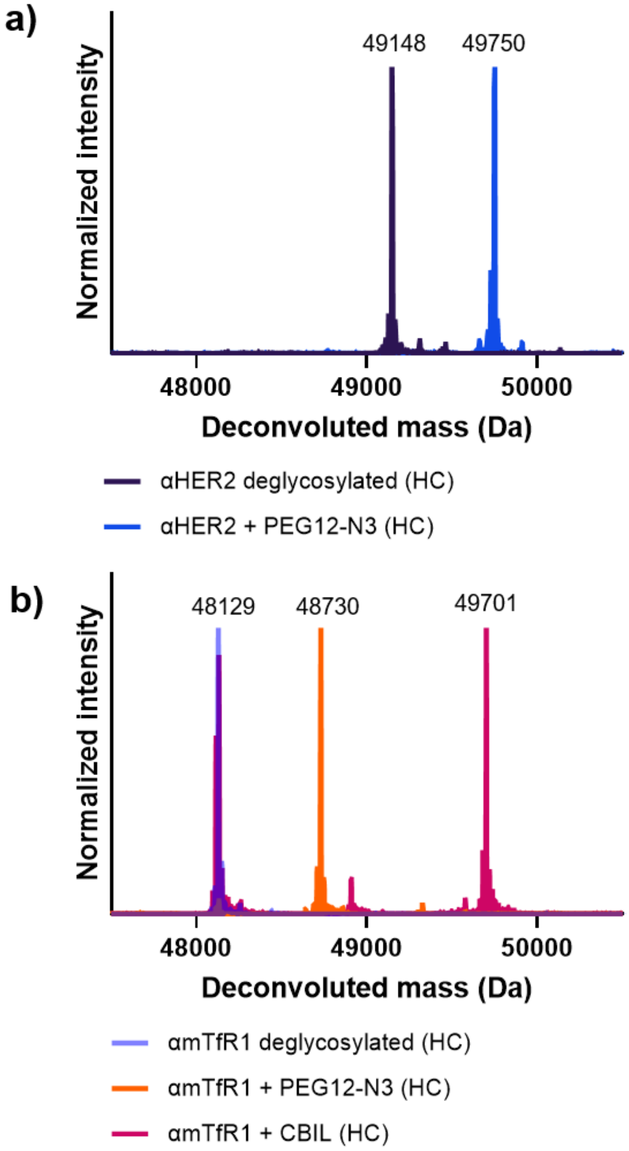

**Figure S11.** Deconvoluted mass spectra of antibody heavy chain (HC) fragments. (a) αHER2 after deglycosylation and modification with PEG12-$N_3$ linker. (b) αmTfR1 after deglysoculation and modification with two different linkers.

#### 5. Gel Electrophoresis methods

##### 5.1 SDS PAGE

SDS-PAGE was performed using 4–12% NuPAGE Bis-Tris precast gels (Invitrogen) in 1× MES running buffer at 200 V for 35 min. Gels were stained using the rapid staining protocol with SimplyBlue SafeStain (Invitrogen). . Gels were visualized using a ChemiDoc imaging system (Bio-Rad).

##### 5.2 Native agarose

Native agarose gel electrophoresis was performed using 2% agarose gels containing GelRed nucleic acid stain. Gels were run in 1× TBE buffer at 100 V for 45 min. Gels were visualized using a ChemiDoc imaging system (Bio-Rad).

#### 6. Confocal microscopy

C2C12 cells were seeded on glass coverslips in 24-well plates ($1 \times 10^5$ cells/coverslip) and allowed to adhere overnight. Cells were incubated with 0.3 µg/mL Cy5-labeled αHER2 AOC or αmTfR1 AOC for 5, 30, or 60 min before fixation with 4% paraformaldehyde. Following permeabilization with 0.2% Triton X-100 in PBS and blocking with 1% BSA/0.1% Triton X-100 in PBS, cells were stained with Alexa Fluor 555-phalloidin (#U0289, Fisher Scientific) and Hoechst 33342 (#62249, Invitrogen; 1:2000). Coverslips were mounted using Fluorescence Mounting Medium (#S302380-2, Agilent Technologies).

Images were acquired using a Nikon Ti2-AX microscope equipped with the Nikon NSPARC super-resolution module and a Plan Apo λD 60×/1.42 NA oil-immersion objective. Samples were excited with 405, 561, and 638 nm laser lines. All samples within an experiment were imaged using identical laser power, detector settings, acquisition parameters, and image reconstruction settings.

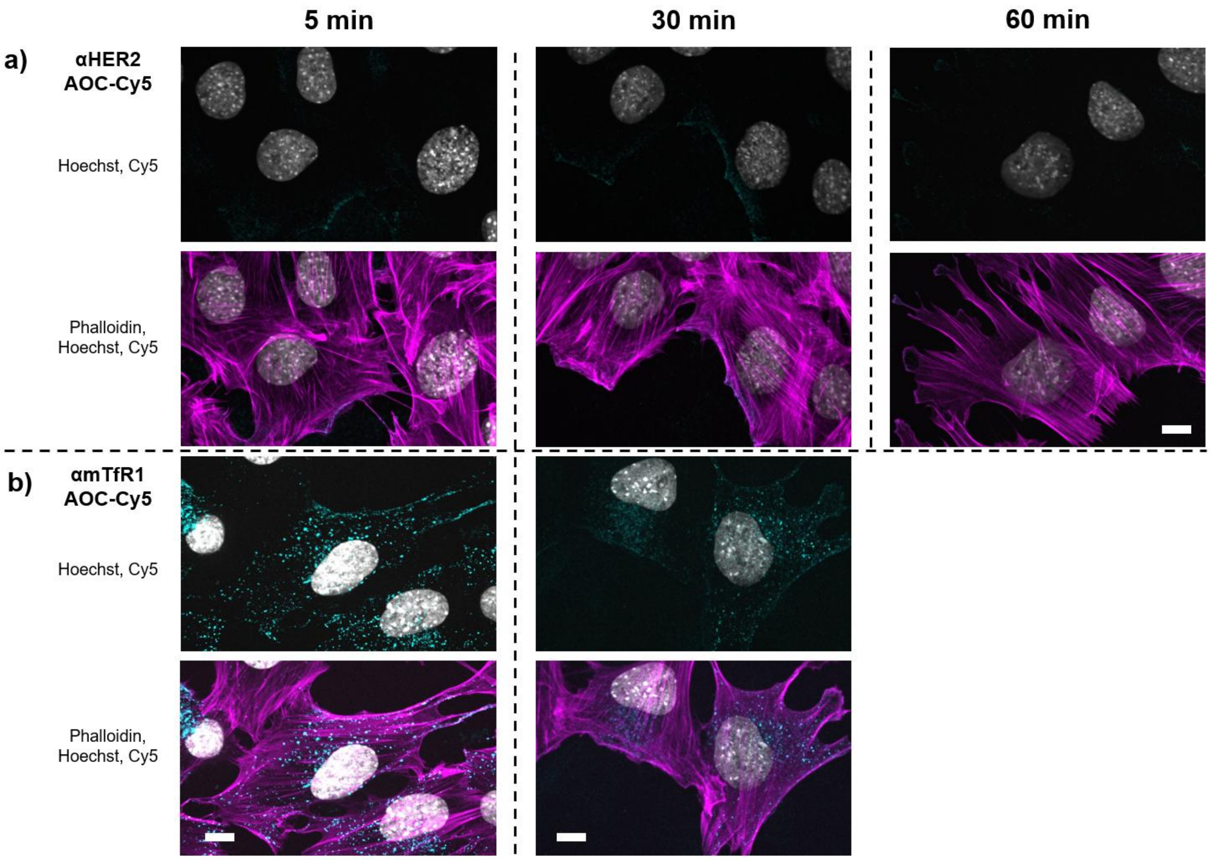

**Figure S12.** A time-course confocal study to investigate internalization of Cy5 labelled AOCs. (a) C2C12 cells treated with Cy5 labelled αHER2 AOC control or (b) αmTfR1 AOC were stained with phalloidin to visualize F-actin and Hoechst to visualize nuclei. Cells were imaged after either 5, 30 and 60 min incubations with respective AOCs. Scale bars = 10 µm.

#### 7. In vitro evaluation of siRNA and model AOC

Neuro-2a cells (N2a) were treated with the indicated siRNA and AOCs at high (400 nM) or low (200 nM) concentrations, with respect to siRNA. Where gymnotic delivery was not assessed, cells were electroporated or transfected with RNAiMAX and analyzed 48 h post-treatment. *Sod1* mRNA levels were quantified by TaqMan qPCR, normalized to Gapdh expression, and expressed relative to untreated controls.

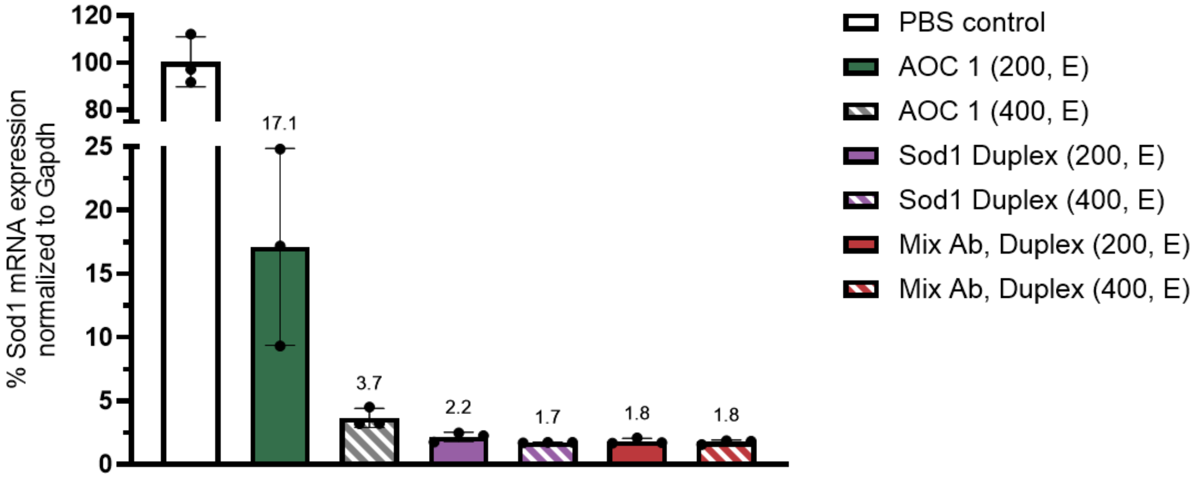

**Figure S13.** *Sod1* knockdown in N2a cells following electroporation (E) of siRNA or AOCs, normalized to the vehicle control. High (400 nM) or low (200 nM) concentrations, with respect to siRNA.

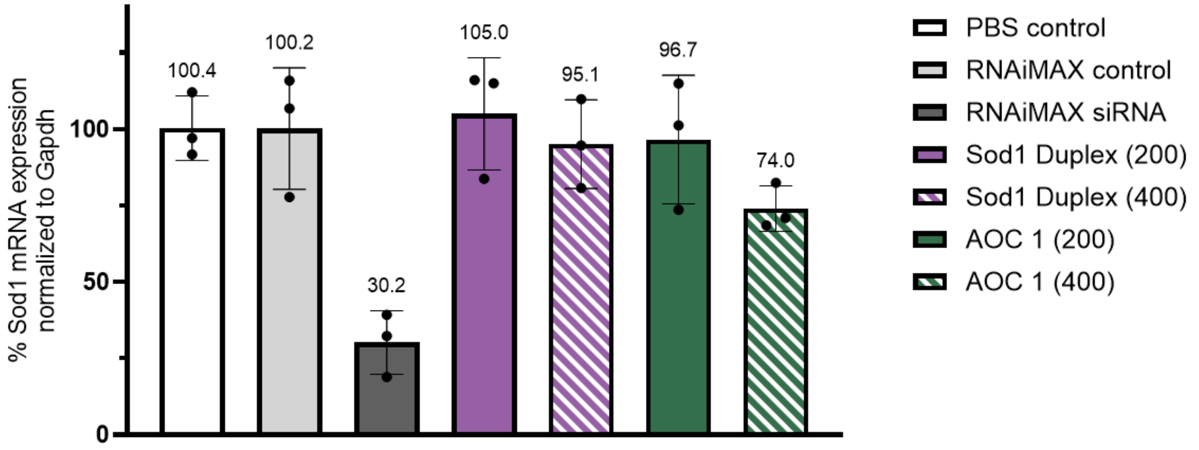

**Figure S14.** *Sod1* knockdown in N2a cells following gymnotic delivery of siRNA or AOCs, normalized to the vehicle control. High (400 nM) or low (200 nM) concentrations, with respect to siRNA.

#### 8. Animal Experiments

Animal experiments were approved by the Swedish Animal Ethics Committee in Linköping, Sweden (permit number 13849-2020; 14772-2023) under the supervision of the Swedish Board of Agriculture (Jordbruksverket) and conducted in accordance with national legislation and EU Directive 2010/63/EU for animal experimentation. Experimental procedures were designed to minimize animal suffering and the number of animals used. Animals were euthanized using approved humane methods in accordance with institutional and national guidelines.

NMRI mice (female, 4-5 weeks old or 20-25 g) were obtained from Janvier Labs. Animals were housed at the Preclinical Laboratory (PKL), Novum, Karolinska University Hospital, Huddinge, under specific pathogen-free conditions compliant with national animal welfare legislation. Animals were acclimatized for at least 7 days following arrival from the supplier. Mice were group-housed in individually ventilated cages

(IVC; maximum five adult animals per cage) with wood-chip bedding and environmental enrichment including nesting material, gnawing sticks, and shelters. Animals had ad libitum access to food and water and were maintained at an ambient temperature of 20–22 °C, 45–55% humidity, and a 12 h light/dark cycle. Animals were monitored daily by trained animal care staff with veterinary supervision available when required.

#### 8.1 In vivo experiments and knockdown analysis by qPCR

Mice were tail-vein injected with AOCs at 1 mg/kg, with respect to siRNA. Seven days post-injection mice were euthanized, and tissues were harvested and stored at -80C until further processing.

Tissues were homogenized in TRI Reagent® (T9424, Sigma-Aldrich) on TissueLyser II (QIAGEN) with stainless steel beads (heart) or on gentleMACS™ Dissociator (Miltenyi Biotec) using program RNA_02_01 (liver). Total RNA was extracted according to manufacturer's instructions (TRI Reagent®) and reverse-transcribed with High Capacity cDNA Reverse Transcription Kit (43-688-13, Applied Biosystems™). qPCR was performed on a CFX Opus 96 Real-Time PCR Instrument (Bio-Rad) using TaqMan™ Fast Advanced Master Mix (4444557, Applied Biosystems™) and TaqMan® Gene Expression Assays (Sod1 = Mm01344233_g1, Gapdh = Mm99999915_g1). Following the ΔΔCt method, Sod1 expression was analyzed relative to Gapdh, and normalized to the PBS-treated control group.

#### 8.2 Dose optimization trial

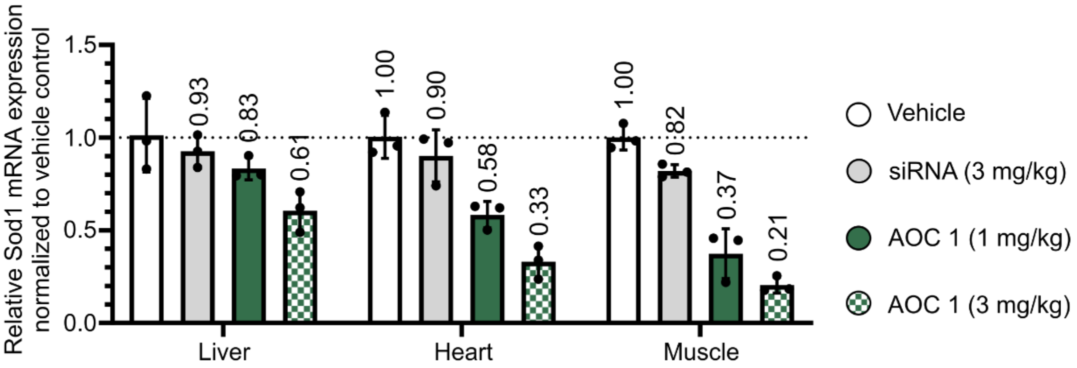

**Figure S15.** *Sod1* knockdown in liver, heart and skeletal muscle tissue normalized to vehicle control for AOCs 7 days post IV injection (1 mg/kg or 3mg/kg). Data are presented as mean ± SD of biological replicates (n = 3).
